# Language exposure and brain myelination in early development

**DOI:** 10.1101/2022.05.31.493945

**Authors:** Laia Fibla, Samuel H. Forbes, Jordan McCarthy, Kate Mee, Vincent Magnotta, Sean Deoni, Donnie Cameron, John P. Spencer

## Abstract

The language environment to which children are exposed has an impact on later language and cognitive abilities as well as on brain development; however, it is unclear how early such impacts emerge. This study investigates the effects of children’s early language environment and socioeconomic status (SES) on brain structure in infancy at both 6 and 30 months of age. We used magnetic resonance imaging (MRI) to quantify concentrations of myelin in specific fiber tracts in the brain. Our central question was whether Language ENvironment Analysis (LENA) measures from in-home recording devices and SES measures of maternal education and family income predicted myelin concentrations over development. Results show relationships between amount of in-home adult input and myelination in the white matter tracts most associated with language. Right hemisphere regions also show an association with SES, with older children from a higher SES background who were exposed to more adult input showing greater concentrations of myelin in language-related areas. We discuss these results in relation with the current literature and implications for future language research and intervention.

**Significance Statement:** This is the first study to look at how brain myelination is impacted by language input and socioeconomic status early in development. We find robust relationships of both factors in language-related brain areas at 30 months of age.

## Introduction

Children’s early language environment is crucial for their emerging language abilities (Hoff & Naigles, 2002). Children exposed to a large quantity of high-quality language input show better language outcomes, enhanced cognitive abilities, and better academic achievement (Rodriguez & Tamis-LeMonda, 2011; Vernon-Feagans et al., 2022; Weizman & Snow, 2001).For example, children whose parents provide both greater quantity and quality of language input – longer utterances, higher grammatical complexity, more vocabulary diversity and more conversational experience – have larger vocabularies (Huttenlocher et al., 2010; Laing & Bergelson, 2019; Rowe, 2012). In addition, children who are exposed to more child-directed speech show faster language processing abilities, becoming more efficient in processing familiar words (Weisleder & Fernald, 2013). Some findings suggest that children might benefit from different aspects of language input at different ages. For instance, overall quantity of input may be more important during the second year of life, diversity in vocabulary input might be more important during the third year of life, and decontextualised language – the use of narrative or explanations – might be particularly beneficial during the fourth year of life (Rowe, 2012).

There are large individual differences in the quantity and quality of language input that children receive from their caregivers. This variation has often been related to parental socio-economic status (SES). SES is an index of a family’s financial resources based on annual income and/or educational achievement based on the number of years of parental education. Some studies report that parents from higher SES backgrounds talk more and use richer language input with their children than parents from lower SES backgrounds (Hart & Risley, 1995; Rowe, 2012). A seminal study showed that, by the time they reach school age, north-American children growing up in higher-SES families hear 30 million more words, on average, than children growing up in lower-SES families (Hart & Risley, 1995). Subsequent research has also shown that, on average, low SES children are exposed to fewer utterances with lower linguistic complexity than higher-SES children (Hoff & Naigles, 2002; Huttenlocher et al., 2007; Huttenlocher et al., 2010; Rowe et al., 2005). Such variations could be due to multiple contextual influences such as high parental stress and economic instability in low SES families as well as different beliefs about infant development and rearing practices (Hoff, 2006). However, recent research has also questioned whether variations in input quantity and quality are strictly associated with SES. A recent study revealed that there was substantial variation in children’s vocabulary environments within each socioeconomic stratum, rather than across different SES groups (Sperry et al., 2019). The authors argued that measuring verbal environments while excluding multiple caregivers and overheard speech produced close to the child – factors which were not taken into account in previous studies (Hart & Risley, 1995) – can underestimate the number of words to which low-income children are exposed to.

Given evidence of a relationship between language input and later academic achievement, what are the mechanisms behind this link? It is possible that higher amounts of language input - particularly words directed to the child, known as ‘child directed speech’ - are associated with increased speech processing ability (Fernald et al., 2013). This mechanism could potentially work in a bi-directional way. Increased child directed speech correlates with speech processing ability already by 18 months (Hurtado et al., 2008) and predicts vocabulary and grammatical development at 24 months (Fernald et al., 2006) as well as language outcomes in elementary school (Marchman & Fernald, 2008). A study with low-SES Spanish-speaking families showed that the amount of speech directed to the child at 19 months – but not overheard speech produced around the child – predicted language processing efficiency and vocabulary size at 24 months (Weisleder & Fernald, 2013). This literature highlights the role of child directed speech as well as the importance of considering multiple caregivers as a source of high quality language input directed to the child.

Recent research has also shown links between language exposure and brain development. A study used home audio recordings to measure children’s language exposure and a story-listening functional MRI task to measure brain activation (Romeo, Leonard et al., 2018). Children between 4 and 6 years of age who had experienced more conversational turns with adults showed greater left inferior frontal activation near Broca’s area in the fMRI task. A mediation analysis showed that activation in this area explained the relation between children’s language exposure and verbal skill. Interestingly, these effects were independent of SES, IQ, and the quantity of adult-child utterances. A second study with the same sample of 4-to-6-year-old children also found a relationship between amount of conversational experience and myelin concentration in white matter tracts most associated with language including the left arcuate fasciculus (AF) that connects Broca’s area with Wernicke’s area as well as the superior longitudinal fasciculus (SLF). Once again, in this study, these relationships were independent of SES and the quantity of adult language input, suggesting that conversational experience has an important impact on brain structure above and beyond SES-related variations in language input. An important question is whether these relationships hold earlier in development when language abilities are first emerging. Another study looked at this question with 6-to 12-month-old infants, showing that characteristics of the home language environment, independent of socioeconomic background, accounted for disparities in early language abilities (Brito et al., 2020). In particular, these authors examined associations between home language input and EEG activity in a socioeconomically diverse sample. They found a positive correlation between measures of socioeconomic status and language input. However, when examining links between language input and brain activity, they found a negative association, with children who heard more adult words in the home demonstrating reduced EEG beta power (13–19 Hz) in parietal regions. Further exploratory analyses revealed a significant interaction between language input and the amount of chaos and disorganization in the home, measured with the Confusion, Hubbub, and Order Scale (CHAOS) scale (Matheny Jr et al., 1995). Children living in high-chaos households who heard more adult words tended to have reduced EEG activity. In contrast, children living in low-chaos homes showed no link between adult input and EEG activity. This study, therefore, revealed complex relationships among language input, SES, the home context, and early brain function. These relationships are particularly difficult to interpret because there is very little research on children’s home language experiences and brain development in infancy. A better understanding of the relationships between language input, environmental factors, and brain development in infancy and early childhood is crucial because it could facilitate the implementation of early intervention programs that boost children’s language abilities during a period when the brain has high plasticity. The present study aims to contribute to this body of research.

The goal of the present study was to investigate the relationships among children’s home language input, SES, and brain myelination in language-related brain regions early in development in a group of 6- and 30-month-old children learning British English. We gathered three types of data: day-long in-home recordings of language experience to infants using the LENA system (Gilkerson et al., 2017), SES information based on maternal education and annual household income, and brain myelination using the mcDESPOT-MRI protocol (Deoni et al., 2012). We obtained individual myelin water fraction maps in all participants (Deoni & Kolind, 2015) and registered them to a common group space to extract myelin concentrations from white matter tracts most associated with language processing and cognitive control: the SLF and the AF.

In an initial analysis, we looked at the relationships between LENA measures (amount of adult words, conversational turns and child vocalizations) and children’s SES based on family income and maternal education. The aim of this first analysis was to see if the LENA output measures followed expected developmental trends across our two age groups including an increase in child vocalizations and conversational turns. Next, we turned to the main focus of the study – we examined how early brain myelination is related to both language exposure and SES by measuring in-home language experience and structural brain development at 6 and 30 months of age. These ages are particularly important because 6-month-old children have high brain plasticity, relatively little brain myelin, and less experience with language. Thus, language input could have a smaller impact on brain structure at this early age. On the contrary, by 30 months of age, most children are able to understand and produce a large number of words (Frank et al., 2017) since after their first birthday, children experience a comprehension boost in which their word learning increases qualitatively (Bergelson, 2020). At 30 months, therefore, language input might have a strong influence on structural brain development, and there might be stronger associations with contextual factors such as SES. A final set of exploratory analyses considered whole-brain myelination in relation to children’s home language experience.

## Materials and Methods

### Experimental Design and Pre-registration

This study was modelled after a recent study looking at the relationships between language input and myelination in the brain (Romeo, Segaran et al., 2018). In our case, we extended these results to a younger population. A priori hypotheses and main analyses were preregistered (see [OSF Pre-registration](https://osf.io/su93b/?view_only=7f7f52fc0fba43a6bbe93a166ac27143)). Our specific hypotheses were: H1) At both 6 months and 30 months, amount of adult input and measures of conversational experience will be positively related to white matter concentrations along fiber tracts known to be involved in language processing and cognitive control: SLF and AF. H2) Measures of conversational experience will be more relevant at older ages as language production increases at 30 months. This should boost the strength of the relationship between conversational turns and white matter concentrations in SLF and AF. We established that our confirmatory analyses would focus on the relationships between language input, SES, and myelin in the AF and the SLF fiber tracks in both hemispheres. We decided to include both hemispheres because at very early ages, brain function is less lateralized than later in development (Deoni et al., 2015). In a set of exploratory analyses, we planned to look at whole-brain myelination since this is the first study to measure language input, SES, and brain myelination early in development.

### Participants

We collected language home input data for 145 children from two age groups: a 6-month-old group (*N* = 83, 38 girls between 4 and 19 months, *M* = 6.94, *SD* = 2.11) and a 30-month-old group (*N* = 64, 35 girls between 28 and 38 months, *M* = 31.15, *SD* = 2.34). A subset of those participants (*N* = 83 children), also had measures of brain myelin using MRI. This subsample included 39 6-month-olds (14 girls; MRIs collected between 5 and 12 months of age) and 44 30-months-olds (23 girls; MRIs collected between 28 and 36 months of age). These participants where white (79), mixed (3) and African (1). All children were native speakers of British English and had no history of premature birth, neurological disorders or developmental delay (see Table 1 for more details).

**Table 1.**
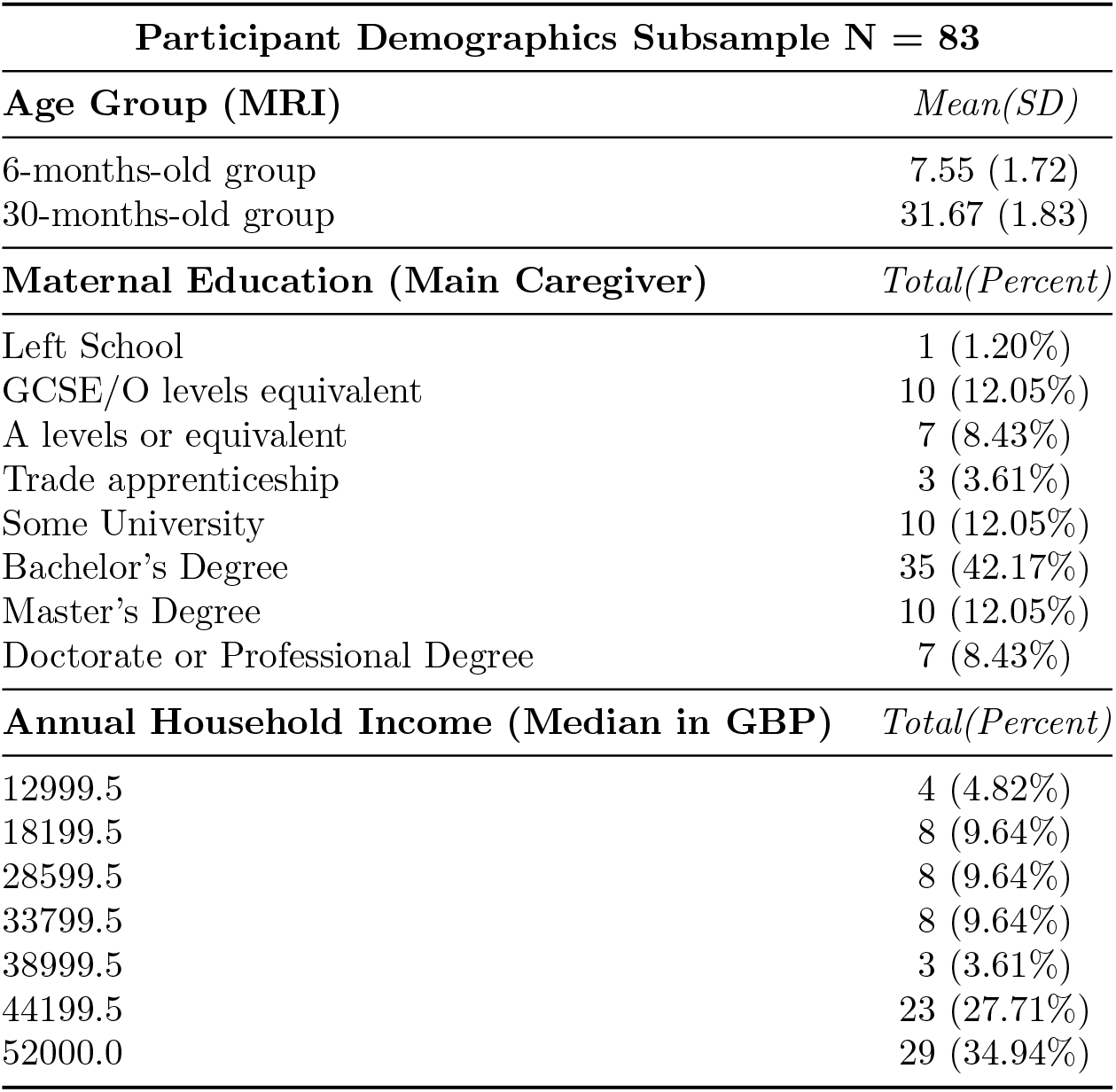
Summary of sample demographics

**Table 2.** Linear Regressions Estimates for the three LENA outcome measures in relation to SEZ

| LENA measure | Term | Estimate | std.Error | p.value |
| --- | --- | --- | --- | --- |
| AWC | (Intercept) | 4958.723 | 163.534 | 0.001 |
|  | AgeGroup | -1351.739 | 327.068 | 0.001 |
|  | SEZ | 405.725 | 192.732 | 0.037 |
|  | Gender | -272.936 | 327.068 | 0.405 |
|  | AgeGroup:SEZ | 160.343 | 385.465 | 0.678 |
|  | AgeGroup:Gender | -121.396 | 654.135 | 0.853 |
|  | SEZ:Gender | 2.909 | 385.465 | 0.994 |
|  | AgeGroup:SEZ:Gender | -722.476 | 770.929 | 0.350 |
| CVC | (Intercept) | 513.786 | 15.543 | 0.001 |
|  | AgeGroup | 340.985 | 31.085 | 0.001 |
|  | SEZ | 30.000 | 18.318 | 0.104 |
|  | Gender | -62.331 | 31.085 | 0.047 |
|  | AgeGroup:SEZ | 17.696 | 36.635 | 0.630 |
|  | AgeGroup:Gender | -29.889 | 62.170 | 0.631 |
|  | SEZ:Gender | -4.195 | 36.635 | 0.909 |
|  | AgeGroup:SEZ:Gender | -12.626 | 73.270 | 0.863 |
| CTC | (Intercept) | 144.438 | 4.628 | 0.001 |
|  | AgeGroup | 75.014 | 9.256 | 0.001 |
|  | SEZ | 8.775 | 5.454 | 0.110 |
|  | Gender | -11.890 | 9.256 | 0.201 |
|  | AgeGroup:SEZ | 3.906 | 10.909 | 0.721 |
|  | AgeGroup:Gender | -15.092 | 18.512 | 0.416 |
|  | SEZ:Gender | -11.440 | 10.909 | 0.296 |
|  | AgeGroup:SEZ:Gender | -16.497 | 21.817 | 0.451 |

*Note.* Results for the three linear models with LENA measure as predicted variable: adult word count (AWC), child vocalizations count (CVC) and conversational turn count (CTC). Fixed effects are displayed including age group (6 or 30 months old), SEZ and child gender.

Six additional children from whom we had both language home input data and brain myelin data could not be included in the final analyses because they exhibited excessive movement in their MRI scans, one additional child was excluded because its family did not provide SES information. All procedures were reviewed and approved by the UK NHS Health Research Authority Ethics committee. Parents signed an informed consent form and received £20 for attending the MRI session. Children received a small toy of their choosing and a t-shirt for participating. The participants from this study are also part of a larger longitudinal project examining the early development of working memory and executive function.

#### 0.1 Socioeconomic Status Measures

We gathered information on the socioeconomic background of each participant and their family using a questionnaire that asked the main two caregivers to provide their level of education as well as the family annual household income. To measure SES, we calculated a composite z-score by averaging z-scores for median annual family income and the main caregiver’s education. In all cases, the main caregiver was the child’s mother and thus, we call this variable maternal education (see Table 1). We refer to this composed score between family income and maternal education as SEZ.

### Language Input Measures (LENA)

The linguistic environment of the child was measured using the Language ENvironment Analysis (LENA) Pro system (Gilkerson et al., 2017). The LENA system is composed of a recorder and associated analysis software. The small recorder can be worn in a vest by the target child at home and it can store up to 16 hours of audio recordings. The LENA software automatically processes the recordings and estimates the number of words spoken by an adult in the child’s vicinity which is referred as adult word count (AWC), the number of vocalizations the target child made or child vocalizations count (CVC), and the number of dyadic conversational turns or conversational turn count (CTC), which is defined as a discrete pair of consecutive adult and child utterances in any order, with no more than 5 seconds of separation. Families took the LENA recorder home on three different days when they did not attend nursery. During those days, the child wore the recorder in a specially constructed vest between 9 and 16 hours (in total we gathered 5712.21 hours of LENA recordings). Each child contributed between 1 and 3 days of recordings. We processed the recordings using the LENA Pro software, which automatically calculated the estimates for each measure (adult words, child vocalizations, and conversational turns). These data were then processed with R (R Core Team, 2021) using a similar approach as in previous literature (Romeo, Segaran et al., 2018). In particular, for each LENA outcome measure and participant, we calculated the total count for each consecutive 60 min across all LENA days, in 5 min increments. For example, we extracted the total amount of adult words that the child was exposed to between 7AM and 8AM, then we calculated the total amount of adult words between 7:05AM and 8:05AM, and so on. We then selected the hour with the highest number of adult words (i.e., the max hour). We used this procedure to extract the hour with the maximum adult word count, the hour with the maximum child vocalizations count, and the hour with the maximum turn count across the several days of home recordings that each participant provided. This maximum measure was used throughout all the analyses reported here.

### Myelin Data Acquisition

The MRI scans where gathered at the *Anonymised* Prior to scanning, children were allowed to fall asleep in a ‘sleepy room’ adjacent to the MRI room. To maximize success, we used these strategies: moved sleeping children into the scanner with minimal disturbance using transportation carts and immobilizers, added a sound-insulating insert to the MR bore (Ultra Barrier, American Micro Industries), used electrodynamic headphones (Ultra Barrier, American Micro Industries), used electrodynamic headphones (MR Confon, Germany), and used customized ‘quiet’ imaging sequences (Deoni et al., 2011). Participants were scanned during natural sleep. Each participant was imaged using a 3T Discovery 750w MRI scanner (GE Healthcare, Milwaukee, WI, USA) equipped with an 8-channel head coil.

### Myelin Data Protocol (mcDESPOT)

For all participants, MRI data was gathered during a period of natural sleep. Myelin content was mapped using a multicomponent driven equilibrium single pulse observation of T_1_ and T_2_ (Deoni et al., 2008). Parameters were as follows: repetition time = 750 ms, echo time = 0.02 ms, inversion time = 650 ms, flip angle = 5^°^, receiver bandwidth = 244 Hz/ voxel, field-of-view = 200 mm x 200 mm, matrix size = 200 × 200, and section thickness = 1 mm. The sequences used as part of the mcDESPOT protocol were: two balanced steady-state free precession (bSSFP) series with phase-cycling increments 0 and 180^°^ to allow for correction of off-resonance artifacts (Deoni, 2011); 8 spoiled gradient echo (SPGR) scans collected over different flip angles; two inversion-recovery SPGR (IR-SPGR) scans for accurate estimation of the B_1_ transmit field. Further, all mcDESPOT data were acquired in pure sagittal or coronal orientation, with a field-of-view adjusted for head size and participant orientation, and a matrix size and section thickness chosen to give consistent isotropic resolution of 1.7 × 1.7 × 1.7 mm^3^. To reduce acoustic noise, these scans were run with reduced gradient amplitudes and slew rates. This resulted in extended scan time. To minimize scan time, mcDESPOT data were acquired with a partial Fourier factor of 0.75 in k_y_ and with an ASSET parallel imaging factor of 1.5. The full protocol lasted less than 45 minutes. A member of the research team was present in the scanner suite to monitor the child at all times.

### Myelin Data Processing

First, the SPGR image with the highest flip angle was selected, and the individual SPGR, IR-SPGR and bSSFP images were all linearly coregistered to that image using *flirt* from FSL (Jenkinson et al., 2002). This accounted for small amounts of motion during the scans. Non-brain tissue and background were then removed from the images. Both the main (B_0_) and transmit (B_1_) magnetic field inhomogeneities were calculated. Myelin water fraction (MWF) maps were then calculated in a voxel-wise manner for every subject using the three-pool model (Deoni et al., 2013). The resulting images were then aligned to a custom template using ANTS (Avants et al., 2011). Core white matter tract masks were used to extract the values for the regions examined, namely the bilateral Arcuate Fasciculus (AF) and the Superior Longitudinal Fasciculus (SLF) so that only voxels contained in these masks were used for analyses. To create the masks, we used a white matter atlas based on the Providence data set (Deoni et al., 2012), except for the AF mask, which was based on a dataset from the University of Manitoba.

#### 0.2 Statistical Analyses

Statistical analyses were conducted using the R software version 4.1.0 (R Core Team, 2021) using the *lm* function. We did three sets of analyses. Within each set, linear regression models followed a basic structure. Analysis 1 looked at relationships among the LENA output measures (number of adult words, conversational turns, and child vocalizations) set as the predicted variable and SEZ set as the predictor variable. The model also included fixed effects of child gender and age group interacting with each other. Analysis 2 (confirmatory) used linear regressions to assess whether language input measures predicted myelination in the SLF and AF. Analysis 3 (exploratory) measured the relationship between language input and other brain tracts that have been related to language in previous literature. The model basic structure used on Analyses 2 and 3 set mean myelin concentration on a specific region as the predicted variable and LENA measure as predictor variable. The models controlled for SEZ and age group set as fixed effects and interacting with each other. Child gender was not included in Analysis 2 and 3 because we did not find consistent effects during Analysis 1, using the LENA data only, and it did not improve model fit for AWC (*χ*^2^(4) = 5537198, *p* = 0.8302), CVC (*χ*^2^(4) = 137812, *p* = 0.3957) or CTC (*χ*^2^(4) = 10097, *p* = 0.4981). In our exploratory analyses, we corrected for multiple comparisons, setting our alpha level for the family-wise error at 0.01. Thus, only *p* values less than 0.01 are considered significant. In all our models, age was included as a categorical variable. This refers to the approximate age when the data were collected. This decision was based on the distribution of age in months, which showed two clusters around 6 and 30 months, and a gap in between. Categorical variables such as age group and child gender were contrast coded, so that one was set as −0.5 and the other set at 0.5.

## Results

### Analysis 1: Language Exposure at Home

Our initial analysis included three linear models, one per LENA measure. The LENA measure was the dependent variable and SEZ, child gender, and age group were set as predictor variables interacting with each other. This means that we assumed that different values on SEZ, gender and age influence each of the LENA measures differently, depending on the values of the other interacting variables.

The linear model predicting the number of adult words showed main effects of SEZ and age group (see Table2). As can be seen in Figure1A, children from families with a higher SEZ score heard more adult words at home that children from families with lower SEZ scores. Moreover, the number of adult words decreased by age – older children heard fewer adult words than younger children. The linear model predicting child vocalizations showed a main effect of age group and a marginal effect of gender. As can be seen in Figure1C, children vocalised more with age as they developed better language skills. Note that boys tended to produce more vocalizations than girls. The linear model predicting amount of conversational turns only showed a main effect of age group with older children producing more turns than younger children (see Figure 1D).

**Figure 1:**
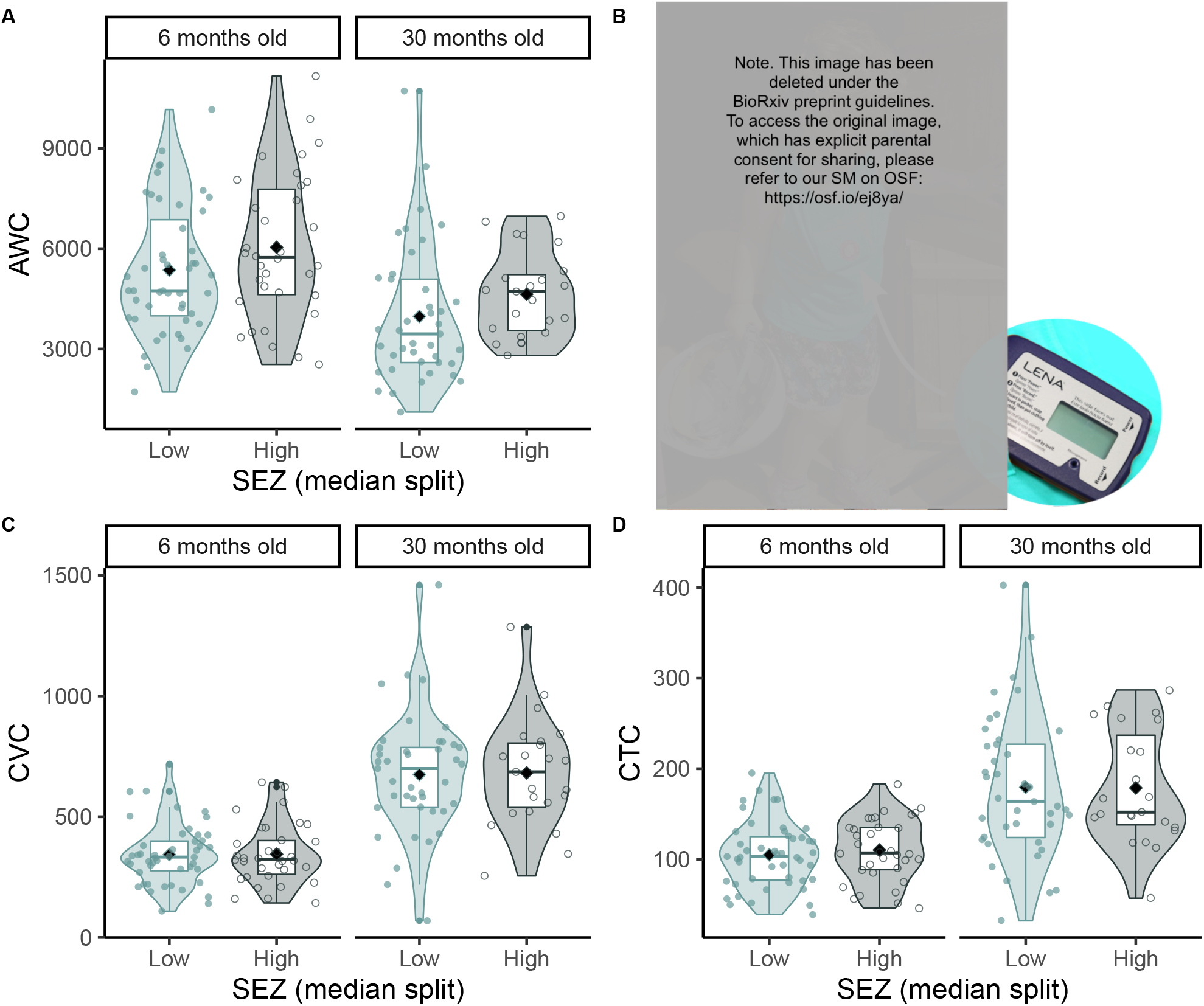
LENA measure showing adult word count (AWC) on panel A, a child wearing the LENA vest with a LENA device inside a pocket on panel B, LENA measure showing child vocalization count (CVC) on panel C and conversational turn count (CTC) on panel D. All graphs are split by age group (6 months old versus 30 months old) and median SEZ (low SEZ in light green and high SEZ in dark green).

### Analysis 2: Language Exposure and Myelin in Language-Related Fiber Tracts (AF and SLF)

Our second set of analyses used linear models to assess whether the three LENA output measures predict myelination in the SLF and AF fiber tracts. We ran linear models with each language exposure measure (adult words, child vocalisations, and conversational turns) predicting mean myelination in the right and left AF and SLF. Models controlled for age group as well as SEZ.

Results of our confirmatory analyses showed positive relationships between the amount of adult input (AWC) and myelination in the AF and SLF (see Table3). As expected, we found positive main effects of age group across both the right and the left AF and SLF, reflecting the increase in brain myelination with age. We also found an interaction between the amount of adult input and age group in these brain regions. In particular, the amount of adult word input was positively associated with the concentration of myelin in the left AF, the right AF, and the right SLF for the 30-month-old group (see darker shaded linear trends in the middle graphs in Figure2). Thus, older children who heard more adult input had more myelinated language-related fiber tracts. At 6 months this relationship was reversed: infants who were exposed to more adult word input had lower myelin concentrations in the regions of interest (see lighter color shading in the middle graphs in Figure2). In the right hemisphere regions (right AF and right SLF; see Table3), we also found an interaction between the number of adult words, age group, and SEZ. This SEZ effect can be seen in Figure2: older children from families with higher SEZ scores, who were exposed to more adult input, showed greater myelin concentrations in these right hemisphere regions.

**Figure 2:**
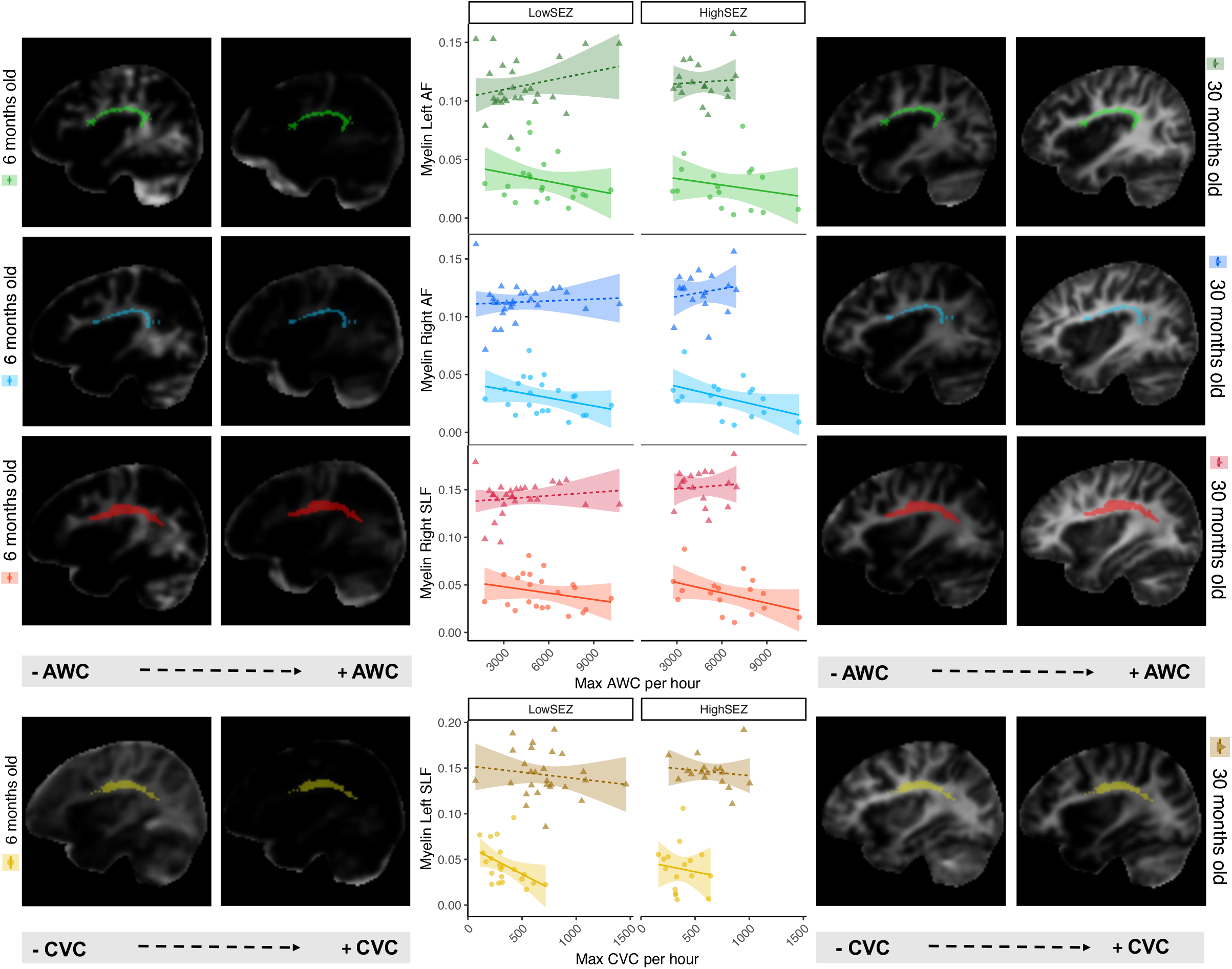
Illustration of the relationships between AWC (adult input) and mean myelin in the left AF (green), right AF (blue) and right SLF (red) as well as CVC (child vocalizations) and mean myelin concentrations on the left SLF (yellow). Dark shading and triangles show data for the 30-months-old group; light shading and dots show data for the 6-months-old group. Scatter plots are divided in low and high SEZ using a median split. Brain images were obtained using MRI scans and show myelin concentration in the brain. The fiber tracts of interest are highlighted on each image using the color scheme from the middle graphs.

**Table 3.** Linear Regression Estimates for AWC predicting myelination in AF and SLF

| Region | Term | Estimate | std.Error | t.statistic | p.value |
| --- | --- | --- | --- | --- | --- |
| Left AF | (Intercept) | 0.073 | 0.006 | 11.270 | 0.001 |
|  | AWC | 0.001 | 0.001 | 0.201 | 0.841 |
|  | AgeGroup | 0.058 | 0.013 | 4.526 | 0.001 |
|  | SEZ | -0.002 | 0.006 | -0.363 | 0.718 |
|  | AWC:AgeGroup | 0.001 | 0.001 | 2.091 | 0.040 |
|  | AWC:SEZ | 0.001 | 0.001 | 0.155 | 0.877 |
|  | AgeGroup:SEZ | -0.003 | 0.013 | -0.255 | 0.799 |
|  | AWC:AgeGroup:SEZ | 0.001 | 0.001 | 0.168 | 0.867 |
| Right AF | (Intercept) | 0.076 | 0.005 | 14.782 | 0.001 |
|  | AWC | -0.001 | 0.001 | -0.177 | 0.860 |
|  | AgeGroup | 0.058 | 0.010 | 5.692 | 0.001 |
|  | SEZ | -0.004 | 0.005 | -0.878 | 0.382 |
|  | AWC:AgeGroup | 0.001 | 0.001 | 2.725 | 0.008 |
|  | AWC:SEZ | 0.001 | 0.001 | 1.317 | 0.192 |
|  | AgeGroup:SEZ | -0.016 | 0.010 | -1.591 | 0.116 |
|  | AWC:AgeGroup:SEZ | 0.001 | 0.001 | 2.045 | 0.044 |
| Left SLF | (Intercept) | 0.096 | 0.007 | 12.988 | 0.001 |
|  | AWC | -0.001 | 0.001 | -0.280 | 0.780 |
|  | AgeGroup | 0.081 | 0.015 | 5.425 | 0.001 |
|  | SEZ | -0.001 | 0.007 | -0.003 | 0.998 |
|  | AWC:AgeGroup | 0.001 | 0.001 | 1.452 | 0.151 |
|  | AWC:SEZ | -0.001 | 0.001 | -0.024 | 0.981 |
|  | AgeGroup:SEZ | -0.010 | 0.015 | -0.643 | 0.522 |
|  | AWC:AgeGroup:SEZ | 0.001 | 0.001 | 0.656 | 0.514 |
| Right SLF | (Intercept) | 0.097 | 0.006 | 16.632 | 0.001 |
|  | AWC | -0.001 | 0.001 | -0.148 | 0.883 |
|  | AgeGroup | 0.073 | 0.012 | 6.328 | 0.001 |
|  | SEZ | -0.002 | 0.006 | -0.356 | 0.723 |
|  | AWC:AgeGroup | 0.001 | 0.001 | 2.717 | 0.008 |
|  | AWC:SEZ | 0.001 | 0.001 | 0.866 | 0.389 |
|  | AgeGroup:SEZ | -0.019 | 0.012 | -1.620 | 0.109 |
|  | AWC:AgeGroup:SEZ | 0.001 | 0.001 | 2.143 | 0.035 |

Note. Fixed effects are displayed including adult word count (AWC), age group (6 or 30 months) and SES z-score (SEZ). Alpha is set at 0.05 (significance level $p < .05$ ).

The second set of linear models examining child vocalizations also showed significant relationships between language experience and myelin concentration. In particular, we found a significant main effect of child vocalizations on brain myelin concentration in the left SLF (see Table4). As can be seen in Figure2, more vocalisations were associated with less myelin in the left SLF. This effect seems to be more pronounced for the 6-month-old group. Our final set of confirmatory models examined brain myelination and turn counts. We did not find any significant relationships between conversational turns and myelin concentrations in the AF and SLF (see Table5).

**Table 4.** Linear Regression Estimates for CVC predicting myelination in AF and SLF

| Region | Term | Estimate | std.Error | t.statistic | p.value |
| --- | --- | --- | --- | --- | --- |
| Left AF | (Intercept) | 0.083 | 0.006 | 13.107 | 0.001 |
|  | CVC | -0.001 | 0.001 | -1.824 | 0.072 |
|  | AgeGroup | 0.081 | 0.013 | 6.408 | 0.001 |
|  | SEZ | -0.003 | 0.009 | -0.320 | 0.750 |
|  | CVC:AgeGroup | 0.001 | 0.001 | 0.802 | 0.425 |
|  | CVC:SEZ | 0.001 | 0.001 | 0.290 | 0.773 |
|  | AgeGroup:SEZ | -0.001 | 0.017 | -0.005 | 0.996 |
|  | CVC:AgeGroup:SEZ | -0.001 | 0.001 | -0.148 | 0.882 |
| Right AF | (Intercept) | 0.074 | 0.005 | 14.214 | 0.001 |
|  | CVC | -0.001 | 0.001 | -0.491 | 0.625 |
|  | AgeGroup | 0.078 | 0.010 | 7.463 | 0.001 |
|  | SEZ | 0.002 | 0.007 | 0.219 | 0.827 |
|  | CVC:AgeGroup | 0.001 | 0.001 | 0.971 | 0.335 |
|  | CVC:SEZ | 0.001 | 0.001 | 0.289 | 0.774 |
|  | AgeGroup:SEZ | 0.021 | 0.014 | 1.445 | 0.153 |
|  | CVC:AgeGroup:SEZ | -0.001 | 0.001 | -1.321 | 0.190 |
| Left SLF | (Intercept) | 0.107 | 0.007 | 15.108 | 0.001 |
|  | CVC | -0.001 | 0.001 | -2.263 | 0.027 |
|  | AgeGroup | 0.094 | 0.014 | 6.679 | 0.001 |
|  | SEZ | -0.006 | 0.010 | -0.587 | 0.559 |
|  | CVC:AgeGroup | 0.001 | 0.001 | 1.245 | 0.217 |
|  | CVC:SEZ | 0.001 | 0.001 | 0.653 | 0.516 |
|  | AgeGroup:SEZ | 0.001 | 0.019 | 0.007 | 0.994 |
|  | CVC:AgeGroup:SEZ | -0.001 | 0.001 | -0.265 | 0.792 |
| Right SLF | (Intercept) | 0.097 | 0.006 | 16.586 | 0.001 |
|  | CVC | -0.001 | 0.001 | -0.934 | 0.353 |
|  | AgeGroup | 0.096 | 0.012 | 8.233 | 0.001 |
|  | SEZ | -0.002 | 0.008 | -0.215 | 0.830 |
|  | CVC:AgeGroup | 0.001 | 0.001 | 1.098 | 0.276 |
|  | CVC:SEZ | 0.001 | 0.001 | 0.911 | 0.365 |
|  | AgeGroup:SEZ | 0.028 | 0.016 | 1.750 | 0.084 |
|  | CVC:AgeGroup:SEZ | -0.001 | 0.001 | -1.731 | 0.087 |

*Note. Fixed effects are displayed including child vocalisation count (CVC), age group (6 or 30 months) and SES z-score (SEZ). Alpha is set at 0.05 (significance level $p < .05$ ).*

**Table 5.** Linear Regression Estimates for CTC predicting myelination in AF and SLF

| Region | Term | Estimate | std.Error | t.statistic | p.value |
| --- | --- | --- | --- | --- | --- |
| Left AF | (Intercept) | 0.079 | 0.007 | 11.417 | 0.001 |
|  | CTC | -0.001 | 0.001 | -0.925 | 0.358 |
|  | AgeGroup | 0.092 | 0.014 | 6.608 | 0.001 |
|  | SEZ | -0.012 | 0.009 | -1.288 | 0.202 |
|  | CTC:AgeGroup | -0.001 | 0.001 | -0.222 | 0.825 |
|  | CTC:SEZ | 0.001 | 0.001 | 1.170 | 0.246 |
|  | AgeGroup:SEZ | 0.006 | 0.019 | 0.340 | 0.735 |
|  | CTC:AgeGroup:SEZ | -0.001 | 0.001 | -0.557 | 0.579 |
| Right AF | (Intercept) | 0.074 | 0.006 | 12.923 | 0.001 |
|  | CTC | -0.001 | 0.001 | -0.374 | 0.709 |
|  | AgeGroup | 0.081 | 0.011 | 7.111 | 0.001 |
|  | SEZ | -0.010 | 0.008 | -1.298 | 0.198 |
|  | CTC:AgeGroup | 0.001 | 0.001 | 0.517 | 0.607 |
|  | CTC:SEZ | 0.001 | 0.001 | 1.425 | 0.158 |
|  | AgeGroup:SEZ | 0.016 | 0.016 | 1.020 | 0.311 |
|  | CTC:AgeGroup:SEZ | -0.001 | 0.001 | -1.088 | 0.280 |
| Left SLF | (Intercept) | 0.102 | 0.008 | 13.102 | 0.001 |
|  | CTC | -0.001 | 0.001 | -1.088 | 0.280 |
|  | AgeGroup | 0.111 | 0.016 | 7.094 | 0.001 |
|  | SEZ | -0.017 | 0.011 | -1.595 | 0.115 |
|  | CTC:AgeGroup | -0.001 | 0.001 | -0.164 | 0.870 |
|  | CTC:SEZ | 0.001 | 0.001 | 1.442 | 0.153 |
|  | AgeGroup:SEZ | 0.002 | 0.021 | 0.090 | 0.928 |
|  | CTC:AgeGroup:SEZ | -0.001 | 0.001 | -0.374 | 0.709 |
| Right SLF | (Intercept) | 0.095 | 0.006 | 14.687 | 0.001 |
|  | CTC | -0.001 | 0.001 | -0.438 | 0.663 |
|  | AgeGroup | 0.099 | 0.013 | 7.663 | 0.001 |
|  | SEZ | -0.012 | 0.009 | -1.388 | 0.169 |
|  | CTC:AgeGroup | 0.001 | 0.001 | 0.598 | 0.552 |
|  | CTC:SEZ | 0.001 | 0.001 | 1.622 | 0.109 |
|  | AgeGroup:SEZ | 0.024 | 0.018 | 1.386 | 0.170 |
|  | CTC:AgeGroup:SEZ | -0.001 | 0.001 | -1.376 | 0.173 |

*Note. Fixed effects are displayed including conversational turn count (CTC), age group (6 or 30 months) and SES z-score (SEZ). Alpha is set at 0.05 (significance level $p < .05$ ).*

Overall, results from the confirmatory analyses showed that the amount of adult language input that children are exposed to is positively associated with brain myelination in the AF and SLF at 30 months. This partially confirms Hypothesis 1; however, we found negative associations between adult word input and brain myelination at 6 months. Our second hypothesis was that conversational experience would positively predict brain myelination at 30 months. This was not supported by our analyses.

### Analysis 3. Language Experience and Overall Brain Myelination

Our last set of analyses aimed to explore the effect of language exposure on overall brain development. We conducted a set of exploratory analyses using a similar set of linear models as in Analysis 2, but now looking in a larger set of brain regions. As in the previous models, we controlled for age group and SEZ. We selected right and left brain regions that have been associated with language development in prior work. We decided to only consider maximum adult input per hour in this set of analyses because the other LENA measures did not show strong relationships to myelin concentrations in our a priori regions of interest.

Most brain areas showed a strong positive age main effect, indicating that myelin concentrations increased with age (see Table6). Recall that we set our alpha level for the family-wise error at 0.01 (*p<*0.01); consequently, only two areas showed significant relationships with adult input and/or SEZ. In particular, myelin concentration in the left Frontal region showed a positive interaction between amount of adult input and age. These results are shown in Figure3. Older children exposed to more adult input showed higher concentrations of myelin in the left frontal regions of the brain. However, this pattern was reversed for younger children, similar to the trends reported for the AF and SLF. Finally, a linear model predicting myelin concentration in the right regions of the Cerebellum showed a positive main effect of SEZ. This indicates that children from families that had higher SEZ scores, had greater myelin concentration in this brain region.

**Figure 3:**
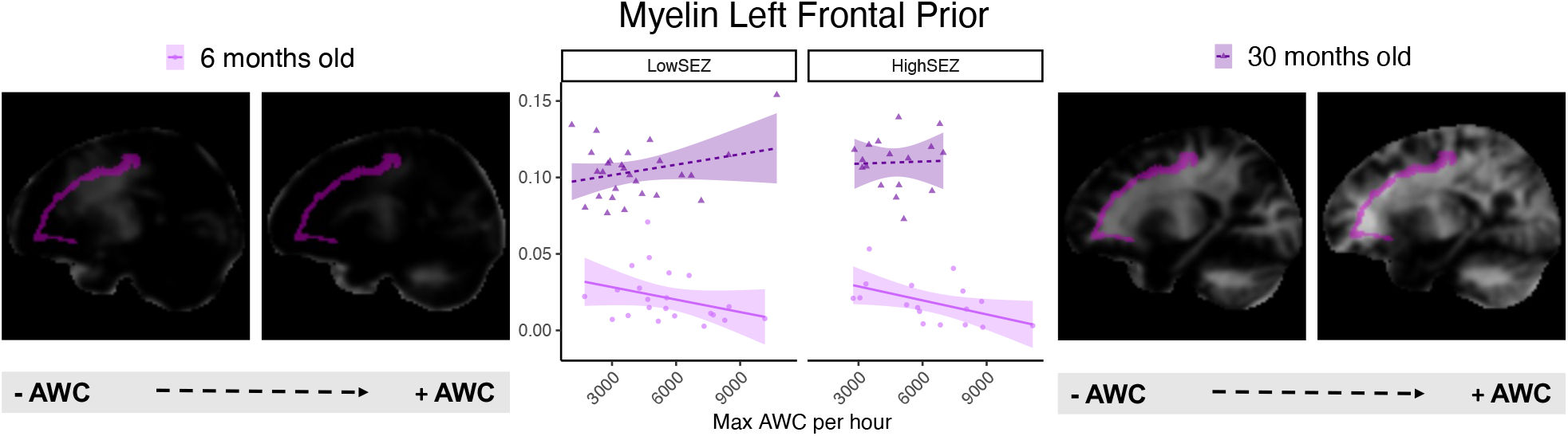
Illustration of the relationships between adult input (AWC) and myelin concentration in the left Frontal Prior. The dark shading and triangles show data from the 30-months-old group, the light shading and dots show data from the 6-months-old group. Middle graph is divided by SEZ using a median split. Brain images were obtained using MRI scans and show myelin concentration in the brain. The left frontal prior fiber tract is highlighted in violet.

**Table 6.**
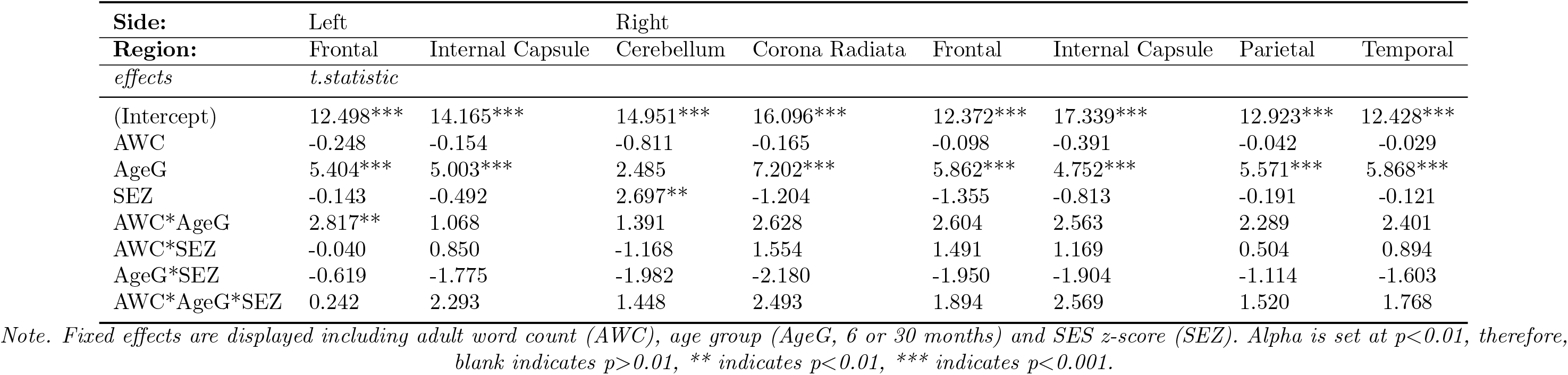
Linear Regression Estimates for AWC predicting myelination in the brain

## Discussion

This study examined the relationship between early language experience and myelination in the brain early in development. We hypothesized that more adult language input and more conversational turns would predict brain myelination in language-related areas, particularly at 30 months of age. Toward that aim, we conducted three analyses with the purpose of quantifying the LENA measures (Analysis 1), confirming or refuting our hypotheses (Analysis 2), and more broadly exploring relationships between children’s language experience and overall brain myelination (Analysis 3). Analysis 1 was used as a primary validation of the LENA system in our sample of participants with the variables of interest. We discovered that the number of adult words was related to children’s SES, with children from higher SES families being exposed to higher amounts of adult input than children from lower SES backgrounds. This finding was somewhat surprising given that our sample was relatively homogeneous with most children coming from middle and high socio-economic backgrounds. This indicates that even small disparities in maternal education and income (combined in our study as a z-score) can have an effect on the amount of adult input children experience early in life. Note, however, that SES was not associated with conversational turns nor with child vocalizations. Analysis 2 was the main focus of this study, quantifying the impact of early language experiences on myelination of the AF and SLF white matter tracts. Results showed that the amount of adult word input was strongly associated with myelin concentration in the AF and SLF. Results from the 30-month-old group followed the pre-registered predicted pattern: more adult input was positively associated with greater myelin concentrations in the left and right AF and the right SLF. Furthermore, this relationship was stronger for higher SES children in the right hemisphere. For the 6-month-old group, we found a surprising negative relationship between adult word counts and myelin concentration. These negative relationships should be interpreted with caution as myelin concentrations are quite low at the age of 6 months and might be susceptible to noise in the MRI data. It is also possible that adult words captured in the vicinity of the infant might contain more overheard speech rather than infant-directed input. Consistent with this, we see fewer conversational turns at earlier ages as parents and infants engage in less babbling or conversational exchanges. Nevertheless, other studies also found negative relationships between home language input and brain activity in children aged 6 to 12 months (Brito et al., 2020). These researchers related this effect to more chaos at home. We did not measure chaos at home; thus, future work will be needed to examine these relationships in more detail.

Our results did not show an effect of conversational turns on AF and SLF myelination as previously reported in 4- to 6-year-olds (Romeo, Segaran et al., 2018). It is possible that conversational turns have an effect on the brain later in development as children learn more language. In fact, studies looking at the relationships between quantity and quality of language input show that children might benefit from different aspects of language input at different time points depending on their language abilities (Rowe, 2012). Early in development, quantity of language input – which in our study was measured by the number of adult words – seems to be more relevant for children’s emerging language skills. In contrast, quality of language input – richness of words, utterance length, and conversational experience – may be more relevant for children at older ages, consistent with effects reported in previous studies (Romeo, Segaran et al., 2018). This would explain why we found that amount of adult input is more predictive of myelin in the AF and SLF at 30 months, while previous research shows that conversational turns are more relevant at 4 to 6 years of age (Romeo, Segaran et al., 2018). Another difference between our findings and prior work is that our results showed effects in both right and left hemispheres for the AF and only the right hemisphere for the SLF. This is consistent with work suggesting that the brain is less lateralized for language early in development, with left areas gaining more specialization for language as children gain language skills. Our results also diverge in that we found SES effects in our sample, with children from higher SES families exposed to more adult words showing higher myelin concentrations in the right AF and right SLF. It is possible that early in development, children are more sensitive to the effects of lower maternal education and income. It is also possible that the amount of language input is more influenced by SES in comparison to conversational turns which was the focus in prior work. Our exploratory analyses looked at possible relationships between language experience and myelin concentrations in a broad range of brain regions. After family-wise correction, results showed relationships between the amount of adult input and myelin concentrations in the left frontal lobe. These results followed the same pattern as our confirmatory analyses: more adult input was associated with more myelin in the left frontal areas of the brain in the 30-months-old group; this relationship was reversed in the 6 month-old group.

Future research could use transcription techniques to analyze the type of speech directed to 6-month-old infants. This would clarify what the LENA AWC measure is capturing at this age. We also note that the current study is one piece of a larger longitudinal study; thus, further analyses at later time points will help disentangle how the 6-month-old findings are related to the findings at 30 months. Ultimately, we hope to understand how language input and brain myelination co-develop within individuals. We also hope to clarify why all of our SES effects were focused in the right hemisphere and how these effects are modulated over development. Previous studies have indicated that SES effects might be less prominent later in development – at least in relation to language. It is possible that SES effects are reduced later in development as other individual differences and cultural factors play out. In summary, our findings suggest that early in development, the amount of home language input from adults is crucial for the development of myelination in language-related brain regions. Moreover, at early ages, myelination seems to be sensitive to small SES disparities. This highlights the need to develop assessment and intervention tools that can boost early language development, particularly when socio-economic disparities are large, helping all children reach their full potential.

## Acknowledgements

This work supported by R01HD083287 from the National Institutes of Health award to JP Spencer as well as by NIH grants R01MH111578 and P50HD103556 to VAM. We thank all the families and children that participated in this study as well as all the DDLab members, staff, students and volunteers that helped with this project.

